# Drug repurposing for therapeutic discovery against human metapneumovirus infection

**DOI:** 10.1101/2022.07.24.501068

**Authors:** Annelies Van Den Bergh, Patrice Guillon, Mark von Itzstein, Benjamin Bailly, Larissa Dirr

## Abstract

Human metapneumovirus (HMPV) is recognised as an important cause of pneumonia in infants, elderly, and immunocompromised individuals worldwide. The absence of an antiviral treatment or vaccine strategy against HMPV infection creates a high burden on the global health care system. Drug repurposing has become increasingly attractive for the treatment of emerging and endemic diseases as it requires less research and development costs than traditional drug discovery. In this study, we developed an *in vitro* medium-throughput screening assay that allows for the identification of novel anti-HMPV drugs candidates. Out of ∼2400 compounds, we identified eleven candidates with a dose-dependent inhibitory activity against HMPV infection. Additionally, we further described the mode of action of five anti-HMPV candidates with low *in vitro* cytotoxicity. Two entry inhibitors, Evans Blue and aurintricarboxylic acid, and three post-entry inhibitors, mycophenolic acid, mycophenolate mofetil, and 2,3,4-trihydroxybenzaldehyde, were identified. Among them, the mycophenolic acid series displayed the highest levels of inhibition, due to the blockade of intracellular guanosine synthesis. Importantly, MPA has significant potential for drug repurposing as inhibitory levels are achieved below the approved human oral dose. Our drug-repurposing strategy proved to be useful for the rapid discovery of novel hit candidates to treat HMPV infection and provide promising novel templates for drug design.

**Highlights:**

- There is currently no treatment against acute HMPV infection.
- We developed a medium-throughput screening for drug repurposing against HMPV infection.
- Evaluation of a large drug library identified 5 candidates with low μM to nM IC_50_ values against HMPV growth.
- The approved drug mycophenolic acid (MPA) is a nM inhibitor of HMPV *in vitro* infection
- Human MPA plasma levels upon oral dosing are 10x higher than MPA IC_50_ against HMPV *in vitro* infection.

## Introduction

Human metapneumovirus (HMPV) is considered as the most common cause of viral respiratory tract infections in infants, together with respiratory syncytial virus (RSV) (Divarathna et al., 2020; van den Hoogen et al., 2003). Specifically, 90% of children worldwide have experienced at least one HMPV infection by the age of five (Davis et al., 2016; Feuillet et al., 2012). HMPV infection is generally associated with upper respiratory tract symptoms such as coughing, sore throat, and fever and typically self-resolves. Severe infections can occur in infants, immunocompromised people, and the elderly, where HMPV spreads to the lower respiratory tract causing pneumonia or bronchiolitis, which may result in hospitalisation (Collins and Karron, 2013; Panda et al., 2014; Schildgen et al., 2011). Recent epidemiological studies suggest that HMPV is responsible for 10-12% of paediatric hospitalisations, associated with a cost of $277 million in the US alone (Davis et al., 2016). In immunocompromised patients with severe HMPV-associated community-acquired pneumonia (CAP), the reported mortality rate is up to 72.7%, higher than the reported severe influenza virus-associated CAP (54.3%) (Choi et al., 2019). Despite the clinical significance of HMPV, there is currently neither vaccine nor antiviral treatment available on the market. Drug repurposing has become an accepted alternative pathway to the traditional slower-paced and expensive drug discovery process. Proven to be highly efficient and relatively low-cost, drug repurposing can lead to the discovery of licensed drugs with well-described safety profiles (Pushpakom et al., 2019). In the present study, we evaluated a commercially available library comprising ∼2400 approved drugs. The compounds that displayed dose-dependent inhibition of HMPV *in vitro* infection were further investigated for their mechanism of action. The identified drugs in our study provide promising candidates for further preclinical evaluation and the basis for the development of novel HMPV therapeutics.

## Materials and Methods

### Cells and viruses

Rhesus monkey kidney epithelial cells (LLC-MK2: ATCC® CCL-7™) were maintained in Opti-MEM™ I Reduced Serum Medium + GlutaMax™ (Thermo Fischer Scientific) supplemented with 2% foetal bovine serum (FBS, Thermo Fischer Scientific) and 1% Anti-Anti 100X (Merck) at 37 °C in a humified atmosphere of 5% CO_2_. Human metapneumovirus genetically modified to express a green fluorescent protein (HMPV-GFP) was obtained from ViraTree (CAN97-83, MPV-GFP1), with the GFP expression used to monitor the infection status. Virus propagation and titration was performed in LLC-MK2 cells using Opti-MEM supplemented with 1.6% TrypLE™ Express (Gibco) and 100 μg/ml CaCl_2_ (OptiMEM_inf_) at 35 °C in a humidified atmosphere of 5% CO_2_, as previously described (Bailly et al., 2021) and detailed in the supplementary information.

### Drug library and compounds

The open access drug library was purchased from Compounds Australia (Griffith University, Australia) and used for the medium-throughput screening (MTS) and dose-response studies. The library includes MicroSource Spectrum FDA, Selleck Kinase, and Epigenetics collections and comprises of 2394 (∼2400) unique structures. The collection was provided in 96-well plates, with each compound diluted to 5 mM in 100% DMSO, in 1 μL. Plates were stored at - 20 °C until use. Compounds selected for follow-up studies were purchased individually. The compounds 2,3,4-trihydroxybenzaldehyde (260843-5G), aurintricarboxylic acid (A1895-5G), patulin (P1639-5MG), mycophenolate mofetil (ML0284-50MG), mycophenolic acid (M3536– 50MG), hexylresorcinol (209465-25G), triclosan (72779-5G) and Evans Blue (E2129) were purchased from Sigma Aldrich. Iretol (HY-13938-50MG) was purchased from Focus Bioscience.

### Medium-throughput screening (MTS) assay

One day before infection, LLC-MK2 cells were seeded at a density of 800 cells/well, in 20 μL, in black, clear-bottom 384 well plates (Corning CellBIND® microplate, Sigma Aldrich). The 5 mM drug stocks were thawed on ice and diluted to 500 μM with dH_2_O, and 5 μL of each compound was added in duplicate to its respective wells. The positive and negative controls wells were supplemented with 5 μL of 10% DMSO-dH_2_O solution. After 15 min of compound contact with cells, 25 μL of a HMPV dilution in OptiMEM_inf_ was added to yield final drug and virus concentrations of 50 μM (1% DMSO) and 500 FFU HMPV per well (including the positive controls), respectively. The negative wells were mock-infected with 25 μL OptiMEM_inf_. The plates were incubated for 1 h at 4 °C to allow synchronised virus binding to cells, followed by 72 h at 35 °C, 5% CO_2_. The MTS assay was validated using the drugs ribavirin (Wyde et al., 2004), a nucleotide analogue, and suramin (Bailly et al., 2021), a sulfonate-containing drug, known to be potent HMPV inhibitors in this concentration range. Infection was measured by *in situ* ELISA as previously described (Bailly et al., 2021) and detailed in the supplementary information. HMPV growth inhibition percentages were calculated using values normalised to the background as follows:

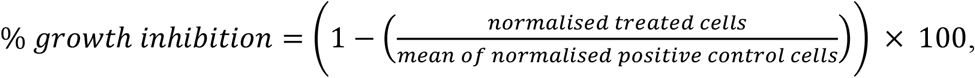

Compounds with obvious cytotoxic effect (80% - 100% cell death) as observed under bright-field microscopy were excluded from further evaluation. The quality of the screening assay was determined by calculating the Z-factor (Zhang et al., 1999) (formula provided in the supplementary information).

### Dose-response studies

Compounds showing more than 50% inhibition of HMPV growth at 50 μM in the MTS were further evaluated for their dose-dependent inhibitory activity against HMPV growth. Their dose-dependent potency was evaluated during all stages of an HMPV infection in the same conditions as the MTS assays. On the day of infection, drug dilutions were prepared on ice in an HMPV OptiMEM_inf_ solution (final 500 FFU/well) starting at 100 μM (final in well), followed by 5-fold serial dilutions and pre-incubated for 15 min on ice. Confluent cell monolayers were incubated with the compound-HMPV solution, in duplicates, for 1 h at 4 °C before transfer to 35 °C for 72 h with 5% CO2. After 3 days, the cells were fixed with 3.7% formaldehyde in PBS and HMPV infection levels measured by *in situ* ELISA, as described in the supplementary information. The dose-response curves were analysed by non-linear regression in GraphPad Prism 8 (GraphPad Software, La Jolla California, USA) to determine the drug concentrations that result in a 50% decrease in infection (IC_50_ values).

### Mode of action studies

The most potent anti-HMPV compounds were further evaluated in four different infection assays to understand their mode of action: (**1**) all-stages, (**2**) binding, (**3**) adsorption, and (**4**) post-adsorption (**Figure 3A**). Confluent LLC-MK2 cells were inoculated with either 500 FFU HMPV per well (**1** and **4**) or 1000 FFU/well (**2** and **3**) for 1 h at 35 °C or 4 °C for the binding (**2**) and adsorption (**3**) assays, respectively. For both the binding (**2**) and adsorption (**3**) assays, 1000 FFU/well of HMPV was used to obtain absorbance signals within an optimal dynamic range to measure differences in infection levels after 1 h inoculation. In all-stages assay (**1**), compounds were present at all stages of infection, while in post-adsorption assay (**4**) compounds were only added after the 1 h HMPV inoculation at 37 °C. In binding assays (**2**), compounds were present during the viral binding only (1 h at 4 °C) and in adsorption assay (**3**) during binding and entry (1 h at 35 °C). For all assays except all-stages (**1**), cells are washed after the 1 h inoculation with OptiMEM_inf_ and further incubated for 72 h at 35 °C and 5% CO_2_. The level of infection was measured by *in situ* ELISA, as described in the supplementary information. The compound concentration resulting in 50% reduction of infection (IC_50_) compared to non-treated infected cells was determined for each compound by non-linear regression of the dose-response curves using GraphPad Prism 8.

### Cytotoxicity

Compound cytotoxicity towards LLC-MK2 cells was determined by using Alamar Blue HS (Thermo Fisher Scientific), in the conditions of dose-response studies and in the absence of HMPV, following the manufacturer’s instructions (see supplementary information).

## Results

### Medium-throughput screening optimisation and validation

To establish the conditions of the medium-throughput screening (MTS), we first determined the HMPV infectious titre required per well to obtain consistent absorbance values after 72 h of infection. In a 384-well plate, we inoculated confluent LLC-MK2 cells with different HMPV titres and measured levels of infection by *in situ* ELISA using an anti-HMPV F primary antibody. As shown in **Figure 1A**, absorbance saturation was reached with 1000 FFU HMPV per well. We therefore continued our MTS validation with 500 FFU/well to allow for a better dynamic range, since signal saturation at 1000 FFU/well may limit signal variations related to compound activity. Next, we validated the MTS using two known anti-HMPV compounds, suramin, a sulfonate-containing approved drug, and ribavirin, an approved nucleoside analogue (Bailly et al., 2021; Wyde et al., 2004). The IC_50_ values for suramin (26.21 μM) and ribavirin (51.05 μM) using our MTS were comparable to the previously reported IC_50_ values of 30.3 μM and 16.38 μM, respectively (**Figure 1B)** (Bailly et al., 2021; Wyde et al., 2004). The assay quality was evaluated using the Z-factor, with a calculated score of 0.87 which is well within the optimal range of 0.5-1 (Zhang et al., 1999). Overall, we concluded that the proposed MTS was suitable for validation of drug libraries against *in vitro* HMPV infection.

**Figure 1.**
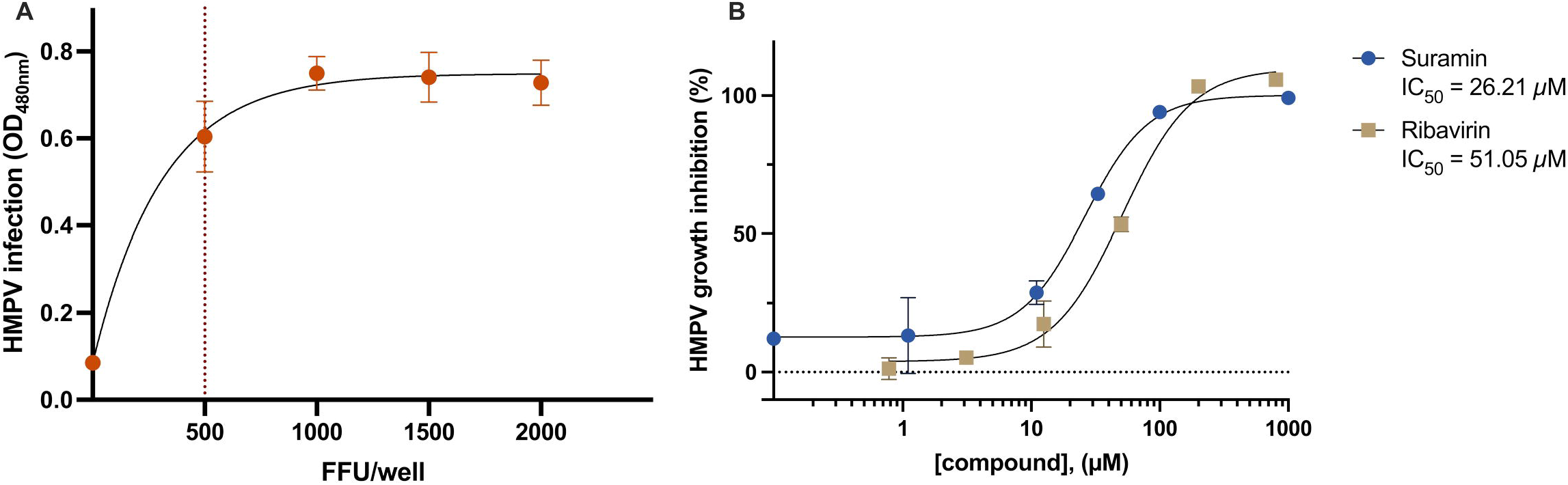
Optimisation and validation of the medium-throughput screening (MTS) assay. (A) Evaluation of the HMPV titre required to obtain absorbance signals with a favourable dynamic range for detection of HMPV inhibition. A confluent monolayer of LLC-MK2 cells was inoculated with various virus titres in a 384 well-plate, and infection was carried out for 3 days. (B) Two known anti-HMPV inhibitors, suramin and ribavirin, were used to evaluate the MTS assay in a dose-dependent manner. HMPV infection levels were measured by *in situ* ELISA using an anti-HMPV F primary antibody. Data points represent the mean ± SD from technical triplicate wells (*n* = 1).

### Medium-throughput screening of an approved drug library

MTS was used to evaluate ∼2400 approved drugs for their potency against HMPV growth *in vitro*. The library, purchased from Compounds Australia, includes drugs from MicroSource Spectrum FDA, Selleck Kinase, and Epigenetics collections, providing a wide range of pharmaceutical indications and targets. Briefly, cell monolayers were pre-treated for 15 min with 50 μM of compound and inoculated with 500 FFU of HMPV. After 72 h, HMPV growth was quantified by *in situ* ELISA. In this MTS, a single concentration of 50 μM was chosen to keep a 1% DMSO concentration. Compounds inhibiting over 50% of infection were selected as hits. A total of 375 compounds showed obvious cytotoxicity resulting in a total disruption of the cell monolayer, as observed by bright-field microscopy (data not shown). They were excluded from further investigation. A total of 61 hits inhibited HMPV growth by more than 50%, and 29 hits by more than 75% (**Figure 2, Table S1**). Overall, no predisposition of anti-HMPV activity for a specific pharmaceutical class was observed, rather we noted a distribution over several classes including anti-microbials and immunosuppressants. However, the screening results are not absolute due to the inherent cytotoxicity of 375 compounds (16% of the library) that could not be further evaluated for their anti-HMPV activity.

**Figure 2.**
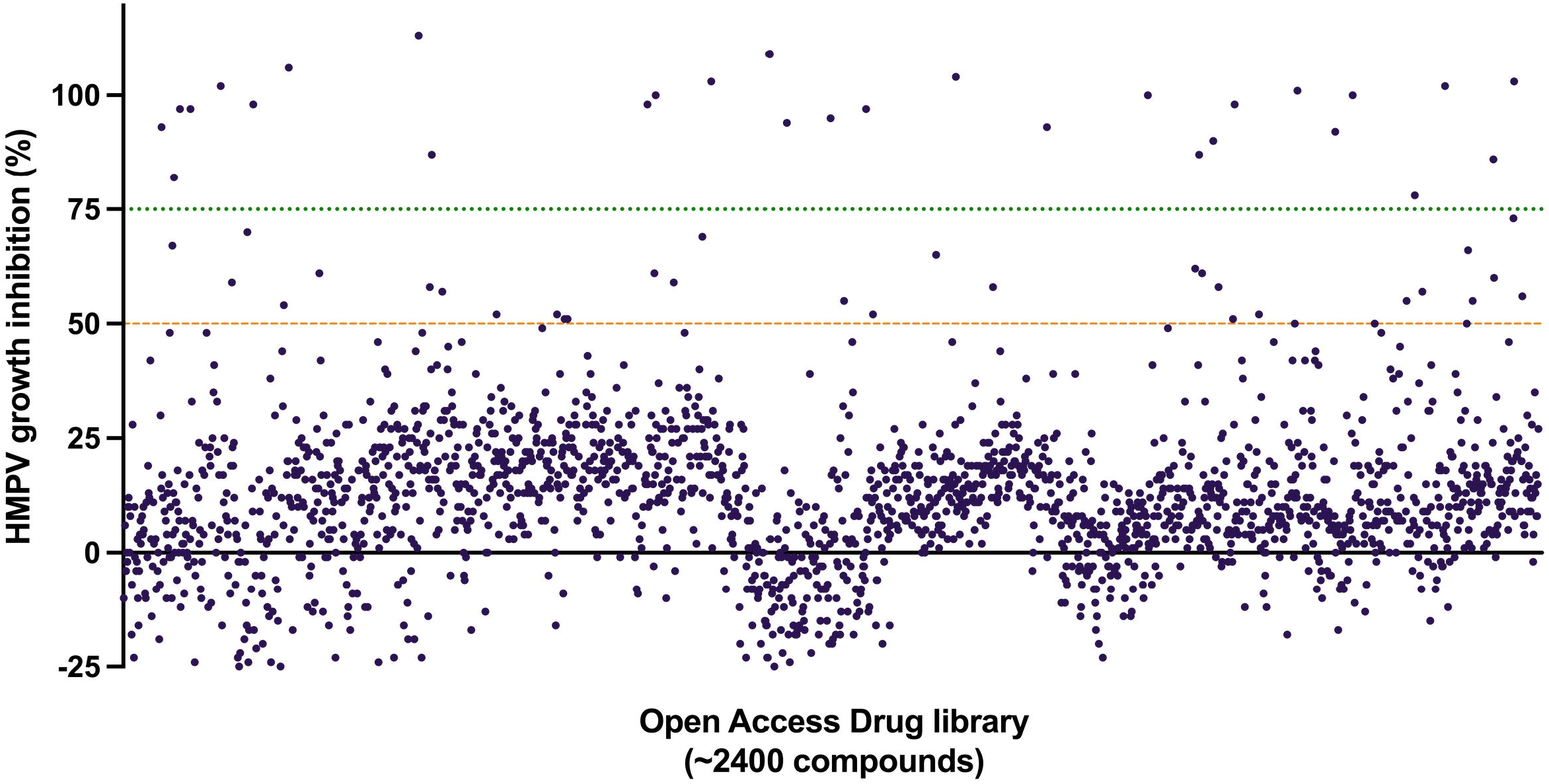
Medium-throughput screening of an approved drug library against HMPV *in vitro* infection. The complete library was evaluated in LLC-MK2 cells, at a compound concentration of 50 μM and compounds were present during all stages of infection using 500 FFU HMPV per well. Compounds that showed more than 50% and 75% inhibition of HMPV growth (presented above the orange and green lines, respectively) were considered as hits for further investigation. Data points represent mean inhibition values of individual compounds, each tested in technical duplicates (*n* = 1), error bars are not shown. Inhibition values of the identified hits are provided in **Table S1**.

### Dose-response evaluation of hits

Next, we determined if the hits selected by MTS inhibited HMPV infection in a dose-dependent manner. For consistency, the hits were sourced from the same provider as the MTS and assays were conducted using the same conditions as the MTS. Among the 61 MTS hits, 11 inhibited HMPV growth in a dose-dependent manner (**Figure S1**) with 50% inhibition concentration (IC_50_) values below 20 μM, as listed in **Table 1**. Moreover, 26 compounds were clearly toxic towards LLC-MK2 cells at a concentration of 100 μM (as observed by bright-field microscopy) and 24 compounds showed a dose-dependent inhibition profile but with high IC_50_ values ≥50 μM. Therefore, they were not further investigated (**Table S2**). IC_50_ values <10 μM were determined for aurintricarboxylic acid (ATA), Evans Blue (EB), patulin, 2,3,4-trihydroxybenzaldehyde (THB), mycophenolic acid (MPA) and mycophenolate mofetil (MPM) (**Table 1**). The selectivity index (*SI = CC*_50_/*IC*_50_) was calculated for each compound. SI values of >20 were obtained for the aforementioned compounds, which demonstrates that the antiviral activities observed are not due to compound cytotoxicity. The compounds may therefore display promising therapeutic windows. Low SI values of 4.5 and 4.8 were calculated for triclosan and hexylresorcinol, respectively, suggesting narrow therapeutic windows. Theaflavin monogallate and irigenol hexaacetate also inhibited HMPV infection in a dose-dependent manner (IC_50_ values <20 μM), although due to limited drug amount availability cytotoxicity assays were not undertaken. Iretol, purchased from another provider (Focus Bioscience), was inactive against HMPV infection and was therefore not evaluated for cytotoxicity. Overall, compounds with IC_50_ values <10 μM against HMPV *in vitro* infection were further investigated for their mode of action due to their significant antiviral potencies (**Table 1**).

**Table 1.**
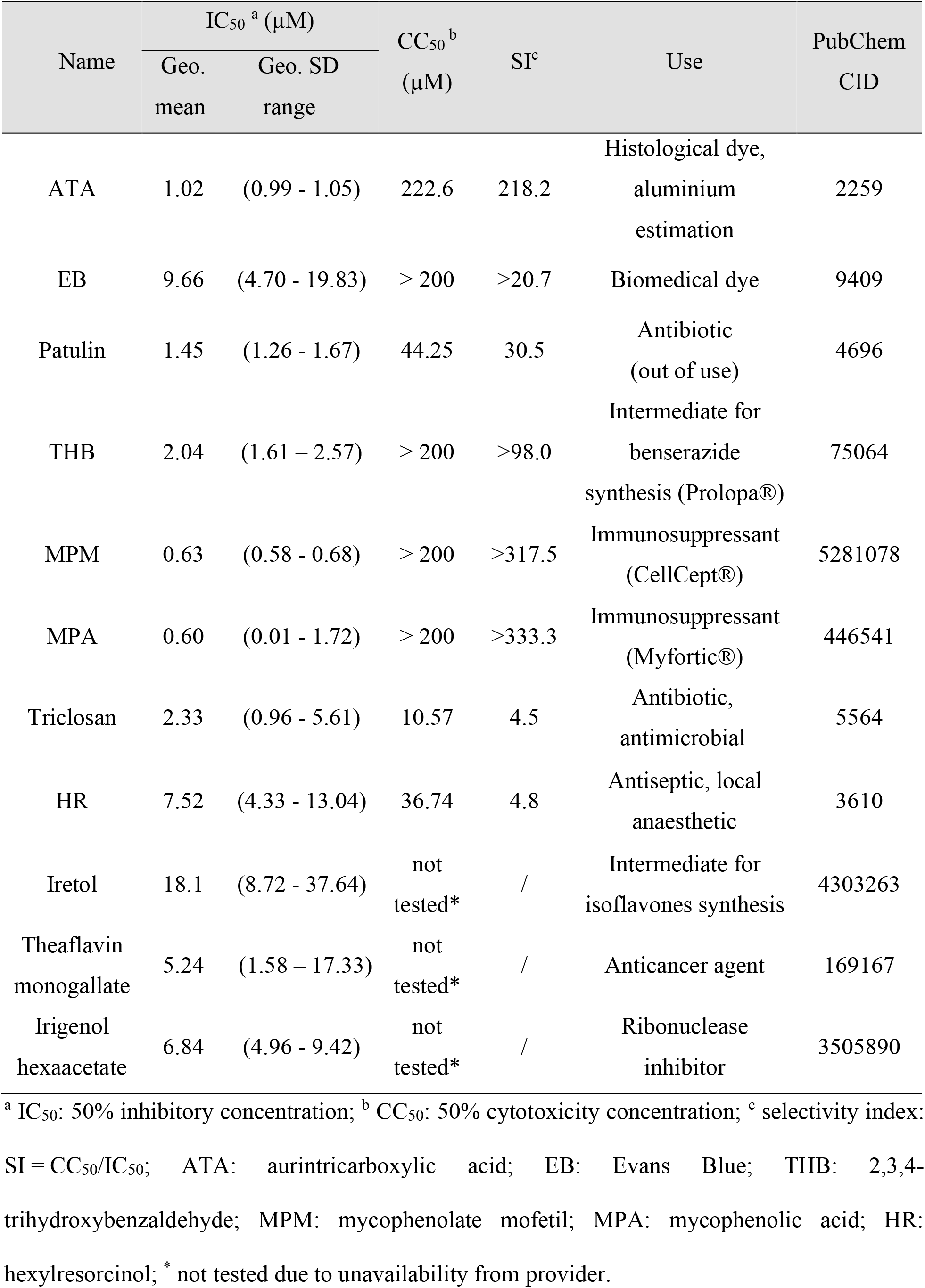
IC_50_ values of selected hits from MTS assay against HMPV infection of LLC-MK2 cells. Infection levels were measured by *in situ* ELISA and cell cytotoxicity was determined using an AlamarBlue assay. The IC_50_ values are calculated fsfigrom two independent experiments performed in technical duplicates and are shown as the geometric mean with their geometric SD range (*n* = 2).

| Name | IC<sub>50</sub><sup>a</sup> (μM) |  | CC<sub>50</sub><sup>b</sup><br>(μM) | SI<sup>c</sup> | Use | PubChem<br>CID |
| --- | --- | --- | --- | --- | --- | --- |
|  | Geo.<br>mean | Geo. SD<br>range |  |  |  |  |
| ATA | 1.02 | (0.99 - 1.05) | 222.6 | 218.2 | Histological dye,<br>aluminium<br>estimation | 2259 |
| EB | 9.66 | (4.70 - 19.83) | > 200 | >20.7 | Biomedical dye | 9409 |
| Patulin | 1.45 | (1.26 - 1.67) | 44.25 | 30.5 | Antibiotic<br>(out of use) | 4696 |
| THB | 2.04 | (1.61 – 2.57) | > 200 | >98.0 | Intermediate for<br>benserazide<br>synthesis (Prolopa®) | 75064 |
| MPM | 0.63 | (0.58 - 0.68) | > 200 | >317.5 | Immunosuppressant<br>(CellCept®) | 5281078 |
| MPA | 0.60 | (0.01 - 1.72) | > 200 | >333.3 | Immunosuppressant<br>(Myfortic®) | 446541 |
| Triclosan | 2.33 | (0.96 - 5.61) | 10.57 | 4.5 | Antibiotic,<br>antimicrobial | 5564 |
| HR | 7.52 | (4.33 - 13.04) | 36.74 | 4.8 | Antiseptic, local<br>anaesthetic | 3610 |
| Iretol | 18.1 | (8.72 - 37.64) | not<br>tested* | / | Intermediate for<br>isoflavones synthesis | 4303263 |
| Theaflavin<br>monogallate | 5.24 | (1.58 – 17.33) | not<br>tested* | / | Anticancer agent | 169167 |
| Irigenol<br>hexaacetate | 6.84 | (4.96 - 9.42) | not<br>tested* | / | Ribonuclease<br>inhibitor | 3505890 |

<sup>a</sup> IC<sub>50</sub>: 50% inhibitory concentration; <sup>b</sup> CC<sub>50</sub>: 50% cytotoxicity concentration; <sup>c</sup> selectivity index: SI = CC<sub>50</sub>/IC<sub>50</sub>; ATA: aurintricarboxylic acid; EB: Evans Blue; THB: 2,3,4-trihydroxybenzaldehyde; MPM: mycophenolate mofetil; MPA: mycophenolic acid; HR: hexylresorcinol; \* not tested due to unavailability from provider.

### Mode of action studies

HMPV treatment can be accomplished by inhibition of either virus binding/entry to cells or replication/spread. To investigate the mode of action of the selected compounds (**Table 1**), they were evaluated in four different assays (**Figure 3A**). During all-stages (**1**) inhibition assays, compound and virus inoculum were present for the 72 h infection period. For binding **(2)** and adsorption (**3**) inhibition studies, cells were inoculated with a mixture of compound and virus for 1 h at 4 °C (binding but no entry) or 35 °C (binding and entry), respectively. For post-adsorption (**4**) inhibition assays, cells were treated post-virus binding/entry, after removing the inoculum. Suramin, previously described as an entry blocker of HMPV infection (Bailly et al., 2021), and ribavirin, a known inhibitor of HMPV genome replication (Wyde et al., 2003), were used as controls. From **Figure 3B**, we can observe that the respective mode of action of the two controls is reflected by their IC_50_ values in the different assays. Ribavirin inhibits HMPV replication after its entry into the cell cytoplasm which only occurs in all-stages (**1**), post-adsorption (**2**), and minimally in the adsorption (**3**) assays. Therefore, a strong antiviral effect for ribavirin was observed at all-stages (**1**, IC_50_ = 40.52 μM) and in post-adsorption (**4**, IC_50_ = 35.15 μM) assays. In contrast, in the binding (**2**) and adsorption (**3**) assays ribavirin is not or not completely able to enter the cell, resulting in higher IC_50_ values. Suramin is known to compete with cellular receptors and prevent HMPV binding to host cells. As expected, we observe similar IC_50_ values for suramin across all four assays as inhibition of HMPV entry can occur at all-stages (**1**, IC_50_ = 37.47 μM), binding (**2**, IC_50_ = 22.61 μM), and adsorption (**3**, IC_50_ = 47.31 μM). A slight increase in the IC_50_ value was noted in the post-adsorption (**4**) assay since initial HMPV binding has occurred prior to suramin addition which only inhibits binding of progeny virus (**Figure 3A**). The evaluation of the compounds outlined in **Table 1** allowed their classification into two categories, entry and post-entry inhibitors, based on their similar inhibition pattern when compared to suramin and ribavirin, respectively (**Figure 3B** and **Table S3**). Mycophenolic acid (MPA), mycophenolate mofetil (MPM) and 2,3,4-trihydroxybenzaldehyde (THB) displayed an inhibition pattern very similar to that of ribavirin. Strong inhibition of HMPV growth by MPA and MPM is observed in all-stages (**1**) and post-adsorption (**4**) but not in the binding (**2**) and adsorption (**3**) assays. However, THB proved to be less potent than ribavirin at all-stages (**1**) and post-adsorption (**4**). While, at all-stages (**1**) and in post-adsorption (**4**) assays, MPA (IC_50_ = 0.21 μM and 0.18 μM, respectively) and MPM (IC_50_ = 0.22 μM and 0.18 μM, respectively) have a 2-log improvement of IC_50_ values compared to ribavirin. The similarity in the IC_50_ values between MPA and MPM are expected as MPM is converted into MPA by intracellular esterases. We therefore infer that MPA and MPM block HMPV replication post-entry. Aurintricarboxylic acid (ATA) and Evans Blue (EB) are classified as entry inhibitors since their inhibition pattern is similar to that of suramin, with their IC_50_ values in the same order of magnitudes across all assays. Moreover, a 1-log improvement of the IC_50_ values for both ATA and EB has been noted compared to suramin when tested at all-stages (**1**, IC_50_ = 1.54 μM and 1.89 μM), in binding (**2**, IC_50_ = 2.37 μM and 2.4 μM) and adsorption (**3**, IC_50_ = 2.52 μM and 4.88 μM) inhibition assays, respectively. All three drugs, ATA, EB, and suramin, demonstrated a slight decrease in inhibition potency when tested post-adsorption (**4**). Interestingly, ATA, EB, and suramin are all negatively-charged drugs at physiological pH which potentially facilitates their interaction with the HMPV surface glycoproteins to compete with the host cellular receptors for attachment.

**Figure 3.**
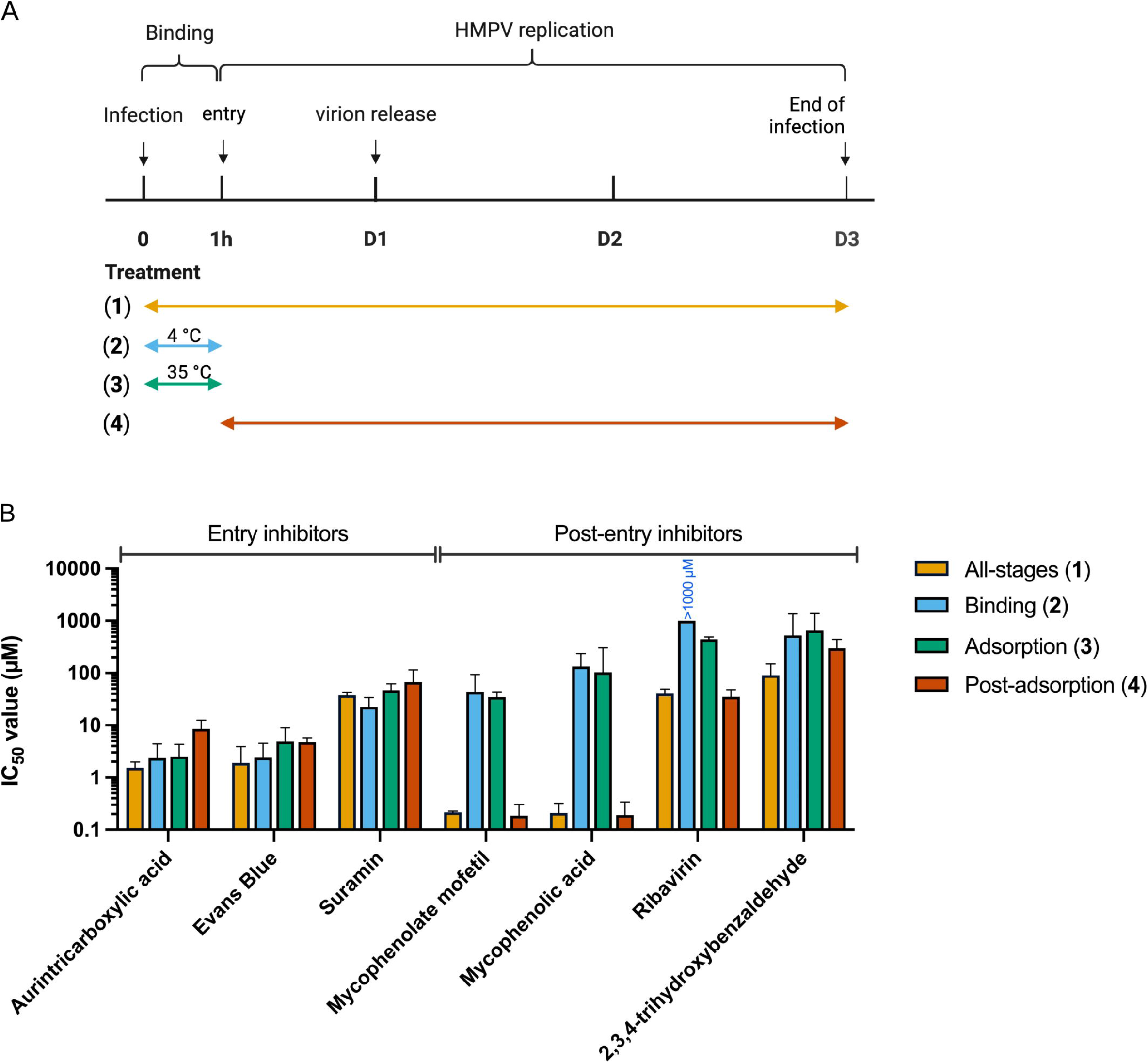
IC_50_ values and mode of action studies of hits against HMPV *in vitro* infection. (A) Time of addition of compounds for the different assays during HMPV infection. (B) The entry inhibitors show binding inhibition activity against HMPV infection, using suramin as a control. The post-entry inhibitors inhibit HMPV replication after internalisation, using ribavirin as a control. Bars represent the geometric mean IC_50_ values from 3 independent experiments and errors bars represent 95% CI (*n* = 3). Inhibition at (1) all-stages, (2) binding, (3) adsorption, and (4) post-adsorption stages. Individual IC_50_ values and their 95% CI for each compound can be found in **Table S3**.

Based on the identification of two groups of inhibitors active at different stages of the HMPV lifecycle, synergy studies were performed. The entry inhibitors ATA and EB were each evaluated in combination with MPA (the active form of the prodrug MPM) as a representative of post-entry inhibitors. Interestingly, no synergistic effect was observed when MPA was combined with either EB or ATA (**Figure S2)**. However, the ZIP-scores of -1 and -1.89, respectively, suggest that an additive effect occurs when MPA is combined with EB or ATA. Finally, three of the eleven dose-response hits identified, patulin, triclosan, and hexylresorcinol, proved to be more toxic towards LLC-MK2 cells (data not shown) than those provided by Compounds Australia, with CC_50_ values of 44.25 μM, 10.57 μM and 36.74 μM, respectively (**Table 1**). Due to their use in consumer products as antimicrobial or local antiseptic, they were not further investigated.

### The anti-HMPV effect of MPA and MPM is caused by depletion of guanosine

Finally, the mode of action of MPA and its prodrug MPM, approved immunosuppressant drugs, was further investigated given their strong anti-HMPV inhibitory activity and high potential for drug repurposing. Both drugs inhibit the inosine-5’-monophosphate dehydrogenase (IMPDH), causing a depletion of the intracellular pool of guanosine, a nucleoside used in the synthesis of DNA and RNA (Allison, 2005; Dasgupta, 2016). Therefore, we investigated the effect of guanosine supplementation on the antiviral effect of MPA. The addition of guanosine during HMPV *in vitro* infection of LLC-MK2 cells with 1 μM MPA [∼ 90% inhibition concentration (IC_90_)] at all-stages (**1**) reversed the antiviral effect of MPA, as shown in **Figure 4**. Moreover, a dose-dependent rescue of the viral infection was observed, with a complete rescue at about 100 μM guanosine supplementation for both MPA and MPM. The antiviral effect of ribavirin, a known nucleoside analogue, was also reversed by guanosine supplementation.

**Figure 4.**
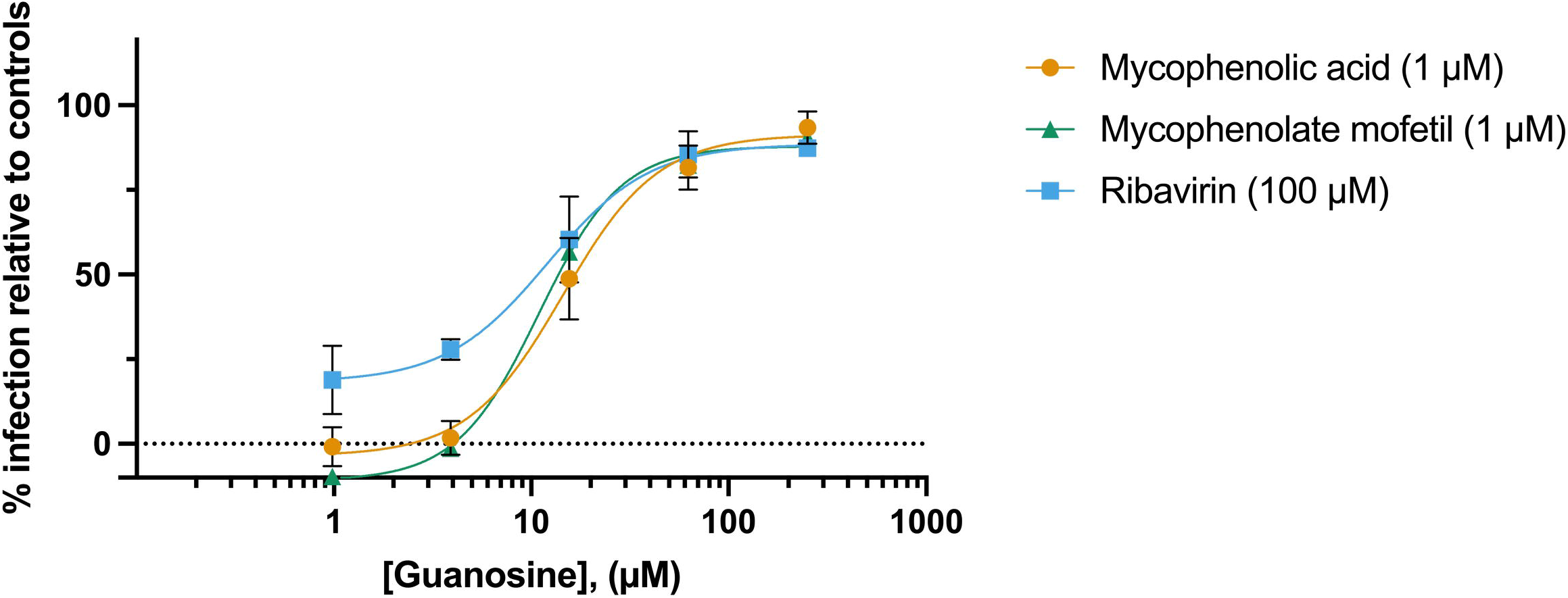
Guanosine supplementation rescues HMPV infection after treatment with mycophenolic acid, mycophenolate mofetil or ribavirin. Infection was performed by mixing 500 FFU HMPV per well, compound, and a dilution series of guanosine overlaid on LLC-MK2 cells for 3 days at 35 °C, 5% CO_2_. The dose-response data points for mycophenolic acid were calculated from three independent repeats performed in triplicates, ± SD (*n* = 3). The data points for ribavirin and mycophenolate mofetil were calculated from two and one independent repeats performed in triplicate (*n* = 2 or 1), respectively.

## Discussion

We report the discovery of novel anti-HMPV compounds identified from a large drug library using an in-house developed and optimised *in vitro* medium-throughput screening (MTS). HMPV is a significant cause of childhood pneumonia worldwide, with no specific therapy available (Choi et al., 2019; van den Hoogen et al., 2003). The need for a readily available drug is therefore critical and the screening of approved drugs provides candidates with established safety profiles. In this study, we used LLC-MK2 cells, a well-established immortalised cell line susceptible to HMPV infection, for compound screening. This cell line allows rapid and consistent virus growth, ideal for MTS (Nao et al., 2019).

Our validated MTS was used to screen ∼2400 approved drugs and we identified 61 drugs with more than 50% inhibition of HMPV growth at a concentration of 50 μM. The hits were further screened for their dose-dependent inhibitory potency against HMPV *in vitro* infection, and 11 drugs demonstrated a high anti-HMPV potency (IC_50_ value < 20 μM). The identified drugs, apart from aurintricarboxylic acid (ATA), were not previously described to possess anti-HMPV properties. A screening by Becker *et al*., 2019 previously evaluated a library of bioactive compounds, natural products, and compounds used in human clinical trials against HMPV *in vitro* infection (Becker et al., 2019). Their screening identified 12 novel compounds, including topoisomerase I/II inhibitors such as ATA, HMG-CoA reductase inhibitors and others. However, their study attributed the anti-HMPV effect of ATA to the antiproliferative mechanism of topoisomerase II inhibitors and did not further investigate its mode of action. The low cytotoxicity of ATA in our study, together with its selectivity index of 218, encouraged us to further investigate its mode of action against HMPV. In this study, we could categorise hits as entry and post-entry inhibitors. The activity of the identified entry inhibitors, ATA and Evans Blue (EB) is likely due to their negative charge at physiologic conditions. We therefore suggest that the observed blockade of HMPV binding is due to charge-based interactions with either the fusion (F) and/or attachment (G) surface glycoproteins. Both surface proteins are involved in viral attachment and entry, but HMPV F was found to be more essential than HMPV G for efficient viral infection (Biacchesi et al., 2004; Cifuentes-Muñoz and Ellis Dutch, 2019). Nonetheless, further studies, e.g. resistance studies, X-ray crystallography, would provide additional characterisation of their respective inhibition profile. Interestingly, ATA was previously described to inhibit *in vitro* infection by other viruses such as Zika virus (IC_50_ = 13.8 μM), Influenza virus (IV) (IC_50_ <10 μM), and Enterovirus 71 (EV71) (IC_50_ = 2.9 μM) (Park et al., 2019; Hung et al., 2009, 2010). ATA is thought to act early in the lifecycle of EV71 and Zika virus and can also inhibit their viral RNA polymerase activity (Park et al., 2019; Hung et al., 2010).The exact mechanism of action, however, is unknown. For IV, ATA has been demonstrated to have a direct inhibitory effect on the neuraminidase activity (IC_50_ value against the IV neuraminidase proteins ranges from 6.3 μg/mL to 16.6 μg/mL and is strain-dependent) (Hashem et al., 2009). The exact nature of the HMPV growth inhibition by ATA cannot be concluded from our results and requires further studies. EB is a well-described medical dye (0.5 – 1% solution) (Yao et al., 2018) and has recently been described as a Severe Acute Respiratory Syndrome Coronavirus 2 (SARS-CoV-2) inhibitor by a proposed dual interaction with the host ACE2 and virus Spike proteins (Day et al., 2021). The observed anti-HMPV effect of EB could possibly be related to interaction of the compound with the HMPV F protein, which is primarily responsible for effective HMPV attachment to the host cell. However, its application as a useful antiviral therapy is unclear due to its non-specific activity and unfavourable pharmacologic profile (Freedman and Johnson, 1969; Yao et al., 2018).

We identified three post-entry inhibitors: mycophenolic acid (MPA), its prodrug mycophenolate mofetil (MPM), and 2,3,4-trihydroxybenzaldehyde. The latter is used as an intermediate to produce the anti-parkinson drug benserazide, used in combination with levodopa in Prolopa^®^. However, its exact mechanism of action against HMPV needs to be further investigated. Both MPA and MPM are approved drugs (Myfortic^®^ and CellCept^®^, respectively) that are administrated as a part of the drug regimen in organ transplants (Dasgupta, 2016). MPA is approved as an immunosuppressant that depletes the intracellular stock of guanosine nucleosides by non-competitive inhibition of the inosine-5′-monophosphate dehydrogenase (IMPDH) (Allison, 2005). Antiviral activity of the mycophenolic acid series has previously been described against Dengue virus (DENV, IC_50_ = 0.4 μM), Middle Eastern Respiratory Syndrome Coronavirus (MERS-CoV) (IC_50_ = 2.87 μM) and SARS-CoV-2 (IC_50_ = 0.87 μM) (Hart et al., 2014; Kato et al., 2020; Takhampunya et al., 2006). In our study, we demonstrated that the anti-HMPV activity of MPA is linked to depletion of the intracellular stock of guanosine. Exogeneous supplementation of guanosine reversed the antiviral effect of MPA, allowing HMPV to replicate and spread. For therapeutic use, the human plasma levels upon oral dosing (bioavailability of MPM is 94%) are approximately 3.5 to 6.5 μM (Dasgupta, 2016), significantly above our estimated IC_50_ value (IC_50_ = 0.21 μM). Therefore, MPA provides an exciting opportunity for immediate clinical use in the treatment of HMPV infection.

Lastly, the entry inhibitors ATA and EB were tested in combination with the post-entry inhibitor MPA for identification of a potential synergistic relationship due to their complementary modes of action. Only a slight additive effect was observed between any of the two drug combinations.

In summary, we have identified five novel drug candidates with potent anti-HMPV *in vitro* activity and low cytotoxicity, from a library of ∼2400 approved drugs, providing new opportunities for the treatment of HMPV infection. A limitation to our study is the use of LLC-MK2 cells, a non-human cell line, and further studies using human-derived cell models are recommended. The compounds identified in this study demonstrate significant potential for repurposing against acute HMPV infection, and provide templates for further novel anti-HMPV drug development.

## Author Contributions

Conceptualization, A.V.D.B. and L.D.; methodology, A.V.D.B., P.G., B.B., and L.D.; analysis, A.V.D.B.; investigation, A.V.D.B.; writing—original draft preparation, A.V.D.B.; writing—review and editing, P.G., M.v.I., B.B., and L.D.; visualization, A.V.D.B.; supervision, P.G., M.v.I., B.B., and L.D.; funding acquisition, M.v.I. and L.D. All authors have read and agreed to the published version of the manuscript.

## Acknowledgments

We thank the team at Compounds Australia at Griffith Institute for Drug Discovery (GRIDD) for the supply of the approved drug library.

## List of Supplementary Tables and Figures

**Table S1: Compounds identified by MTS demonstrating more than 50% HMPV growth inhibition at 50 μM**. The HMPV growth inhibition percentage was calculated from technical duplicates (*n* = 1).

**Table S2: Identified hits with calculated HMPV growth IC**_**50**_ **values above 20 μM and/or obvious cytotoxicity as observed by bright-field microscopy**. The IC_50_ values are calculated from two independent experiments performed in technical duplicate, ± SD, (*n* = 2). ^a^ IC_50_ = 50% inhibitory concentration, *ndr: no dose response.

**Table S2: Individual IC**_**50**_ **values for each hit evaluated in the four different inhibition assays against HMPV infection**. The geometric mean of each IC_50_ value was calculated from at least three independent experiments performed in triplicate and are shown as the geometric mean with their 95% CI, (*n* = 3).

**Figure S1: Dose-dependent inhibition of identified drugs against HMPV-GFP infection of LLC-MK2 cells**. HMPV infectivity was measured by *in situ* ELISA (A) Drugs with IC_50_ values below 10 μM. (B) Drugs with IC50 values above 10 μM. Datapoints represent the mean value from two experimental repeats performed in technical duplicate, ± SD=, (*n* = 2).

**Figure S2. The antiviral potency of drug combinations against HMPV *in vitro* infection using the ZIP method to determine synergy**. The calculations were performed using SyngeryFinder (Zheng et al., 2022) and the synergy score is visualised as the height of a 3D surface. (*n* = 1) (A) The ZIP-score and 3D surface of the synergy score of the combination of mycophenolic acid (MPA) with Evans Blue (EB). (B) The ZIP-score and 3D surface of the synergy score of the combination of MPA and aurintricarboxylic acid (ATA).

