## Supplementary Information for "Drug repurposing for therapeutic discovery against human metapneumovirus infection"

### Supplementary Methods

#### HMPV propagation

Virus culture supernatant was clarified from cell debris 7 days post-infection by centrifugation at 3000 x g for 20 min. The HMPV infectivity titre was determined by focus forming assay. Confluent LLC-MK2 monolayers in a 96-well plate were infected with 10-fold dilutions of HMPV in $OptiMEM_{inf}$ for 1 h at 35 °C and 5% $CO_2$, with gentle agitation every 15 min. The inoculi were aspirated, and cells were overlaid with 0.83% Avicel (FMC BioPolymer) in $OptiMEM_{inf}$. After three days, the cells were fixed with 3.7% formaldehyde (Merck) in PBS for 20 min, followed by immunostaining as described in the section below.

#### Immunostaining

After fixation, cells were permeabilised with 1% IGEPAL CA-630 (Merck) and endogenous peroxidases inactivated with 0.32% $H_2O_2$ (Merck) in PBS for 20 min at 37 °C. Cells were incubated with 0.5 µg/mL mouse monoclonal IgG anti-HMVP F antibody (clone hMPV24, Bio-Rad), followed by a 1:6000 dilution of secondary goat anti-mouse-IgG(H+L)-HRP conjugate (ref# 172-1011, Bio-Rad), each for 1 h at 37 °C. For focus forming assays, 50 µL per well of TrueBlue$^{TM}$ Peroxidase Substrate (Sera Care) was used to obtain blue virus foci. The foci were manually counted, and titres expressed as focus forming unit per millilitre (FFU/ml). For *in situ* ELISA, infection levels were detected by addition of 30 µL (384-well plate) or 50 µL (96-well plate) per well of BD OptEIA™ TMB substrate (BD Biosciences). After 3-5 min, the enzymatic reaction was stopped with 15 µL (384-well plate) or 25 µL (96-well plate) per well of 1 M $H_2SO_4$ and absorbance in each well was measured at 480 nm using an xMark Microplate Spectrophotometer (BioRad). The average absorbance from negative control wells (cells only) was subtracted from the raw data and all values normalised to that of the average from positive control wells (cells and virus), to yield percentages of infection.

**Z-factor calculation**

The quality of the screening assay was determined by calculating the Z-factor using the following formula:

$$\text{Z-factor} = 1 - \frac{3(\sigma_p + \sigma_n)}{|\mu_p - \mu_n|} \quad (1)$$

$\sigma$ = mean

$\mu$ = standard deviation

$p$ = positive control

$n$ = negative control

A screening assay with a Z-factor between 0.5 and 1 is within the qualitative range of an assay (1).

**Cytotoxicity**

After 3 days of incubation with cells and compound, cells were washed and 100 µL Alamar Blue HS in $OptiMEM_{inf}$ was applied to each well. Healthy cells reduce the resazurin from the AlamarBlue solution to resorufin, a red-fluorescent compound, which is detected using a fluorescence spectrophotometer (Tecan Infinite 200 Pro plate reader, Tecan Trading AG, Switzerland) with excitation and emission wavelengths of 560 nm and 590 nm, respectively. The 50% cytotoxic concentration ($CC_{50}$) of compounds was determined by non-linear regression analysis using GraphPad Prism 8.

**Supplementary Tables and Figures**

**Table S1**

| Compound name | HMPV growth inhibition (%) |
|---|---|
| LORNOXICAM | 52% |
| ERYTHRITOL | 65% |
| DICHLOROPHEN | 104% |
| DEOXYSAPPANONE B 7,4-DIMETHYL ETHER | 57% |
| PROTIRELIN | 52% |
| TRICLOSAN | 109% |
| HEXYLRESORCINOL | 106% |
| PRIMULETIN | 61% |
| 4-HYDROXYCHALCONE | 102% |
| MESALAMINE | 95% |
| ADENINE | 58% |
| NORSTICTIC ACID | 87% |
| 4-ACETOXYPHENOL | 82% |
| MYCOPHENOLIC ACID | 87% |
| CEFTRIAXONE SODIUM TRIHYDRATE | 51% |
| 4-METHYLDAPHNETIN | 62% |
| DIGITOXIN | 70% |
| EPICATECHIN MONOGALLATE | 100% |
| IRETOL | 90% |
| GANGALEOIDIN | 66% |
| AMLEXANOX | 51% |
| VIDARABINE | 57% |
| IRIGENOL HEXAACETATE | 98% |
| NORGESTIMATE | 59% |
| PHLORETIN | 67% |
| TRYPTAMINE | 97% |
| OXYTETRACYCLINE | 61% |
| DEOXYSAPPANONE B TRIMETHYL ETHER | 78% |
| EVANS BLUE | 97% |
| LEFLUNOMIDE | 61% |
| THEAFLAVIN MONOGALLATES (TF2b) | 92% |
| RESVERATROL | 51% |
| ISOGINKGETIN | 50% |
| QUINESTROL | 52% |
| ACETOXOLONE | 58% |
| 3-METHYLORSELLINIC ACID | 55% |
| CHICAGO SKY BLUE | 86% |

| | |
|---|---|
| ANTIMONY POTASSIUM TARTRATE TRIHYDRATE | 55% |
| SALINOMYCIN, SODIUM | 58% |
| TERBINAFINE HYDROCHLORIDE | 69% |
| ATOVAQUONE | 59% |
| MYCOPHENOLATE MOFETIL | 100% |
| LAPACHOL | 56% |
| DOBUTAMINE HYDROCHLORIDE | 98% |
| FLUPHENAZINE HYDROCHLORIDE | 113% |
| 2,3,4-TRIHYDROXYBENZALDEHYDE | 100% |
| ETHACRIDINE LACTATE | 93% |
| HEXACHLOROPHENE | 98% |
| α-TOXICAROL (dl) | 93% |
| TETRAC | 60% |
| AZATHIOPRINE | 102% |
| AMLODIPINE BESYLATE | 103% |
| THYMOQUINONE | 73% |
| METHYLENE BLUE | 94% |
| PATULIN | 103% |
| 1-(2-TRIFLUOROMETHYLPHENYL)IMIDAZOLE | 52% |
| DIHYDROERGOTAMINE MESYLATE | 54% |
| PURPUROGALLIN-4-CARBOXYLIC ACID | 55% |
| CHLORHEXIDINE DIHYDROCHLORIDE | 97% |
| AURIN TRICARBOXYLIC ACID | 101% |

**Table S1. Compounds identified by MTS demonstrating more than 50% HMPV growth inhibition at 50 µM.** The HMPV growth inhibition percentage was calculated from technical duplicates (*n* = 1).

**Table S2**

| Compound name | $IC_{50}$ (µM)[a] | Cytotoxicity observed on LLC-MK2 |
|---|---|---|
| LORNOXICAM | >100 | no |
| ERYTHRITOL | ndr* | yes |
| DICHLOROPHEN | ndr | yes |
| DEOXYSAPPANONE B 7,4-DIMETHYL ETHER | >100 | no |
| PROTIRELIN | >100 | no |
| PRIMULETIN | >100 | no |
| 4-HYDROXYCHALCONE | ndr | yes |
| MESALAMINE | 50.4 ± 36.5 | no |
| ADENINE | >100 | no |
| NORSTICTIC ACID | ndr | yes |
| 4-ACETOXYPHENOL | ndr | yes |
| CEFTRIAXONE SODIUM TRIHYDRATE | >100 | no |
| 4-METHYLDAPHNETIN | >100 | no |
| DIGITOXIN | ndr | yes |
| EPICATECHIN MONOGALLATE | 5.4 ± 6.5 | yes |
| GANGALEOIDIN | ndr | yes |
| AMLEXANOX | ndr | yes |
| VIDARABINE | > 100 | no |
| NORGESTIMATE | ndr | yes |
| PHLORETIN | ndr | yes |
| TRYPTAMINE | ndr | yes |
| OXYTETRACYCLINE | 30.8 ± 9.6 | no |
| DEOXYSAPPANONE B TRIMETHYL ETHER | >100 | no |
| LEFLUNOMIDE | 60.4 ± 14.8 | yes |
| RESVERATROL | >100 | no |
| ISOGINKGETIN | ndr | yes |
| QUINESTROL | ndr | yes |
| ACETOXOLONE | >100 | no |
| 3-METHYLORSELLINIC ACID | ndr | yes |
| CHICAGO SKY BLUE | >100 | no |
| ANTIMONY POTASSIUM TARTRATE TRIHYDRATE | ndr | yes |
| SALINOMYCIN, SODIUM | 49.1 ± 15.4 | no |
| TERBINAFINE HYDROCHLORIDE | 74.2 ± 20.8 | no |
| ATOVAQUONE | ndr | no |
| LAPACHOL | 14.2 ± 1.0 | yes |

| | | |
|---|---|---|
| DOBUTAMINE HYDROCHLORIDE | ndr | yes |
| FLUPHENAZINE HYDROCHLORIDE | ndr | yes |
| ETHACRIDINE LACTATE | >100 | no |
| HEXACHLOROPHENE | ndr | yes |
| alpha-TOXICAROL (dl) | ndr | yes |
| TETRAC | ndr | yes |
| AZATHIOPRINE | ndr | yes |
| AMLODIPINE BESYLATE | >100 | yes |
| THYMOQUINONE | ndr | yes |
| METHYLENE BLUE | ndr | yes |
| 1-(2-TRIFLUOROMETHYLPHENYL)IMIDAZOLE | >100 | no |
| DIHYDROERGOTAMINE MESYLATE | >100 | no |
| PURPUROGALLIN-4-CARBOXYLIC ACID | 60.2 ± 3.8 | no |
| CHLORHEXIDINE DIHYDROCHLORIDE | ndr | yes |

**Table S2. Identified hits with calculated HMPV growth $IC_{50}$ values above 20 µM and/or obvious cytotoxicity as observed by bright-field microscopy**. The $IC_{50}$ values are calculated from two independent experiments performed in technical duplicate, ± SD ($n$ = 2). [a] $IC_{50}$ = 50% inhibitory concentration, *ndr: no dose response

### Table S3

| Compound | IC50 (µM) | | | | | | | |
|---|---|---|---|---|---|---|---|---|
| | All stages (1) | | Binding (2) | | Adsorption (3) | | Post-adsorption (4) | |
| | Geo. mean | 95% CI | Geo. mean | 95% CI | Geo. mean | 95% CI | Geo. mean | 95% CI |
| 2,3,4-trihydroxybenzaldehyde | 90.79 | (178.42 - 43.92) | 453.23 | (2428.49 - 84.58) | 612.06 | (1761.08 - 212.72) | 294.31 | (467.05 - 185.46) |
| Mycophenolate mofetil | 0.22 | (0.23 - 0.21) | 39.55 | (165.51 - 9.45) | 34.61 | (45.06 - 26.59) | 0.18 | (0.36 - 0.09) |
| Mycophenolic acid | 0.21 | (0.36 - 0.12) | 130.7 | (268.28 - 63.67) | 102.08 | (723.88 - 14.4) | 0.18 | (0.43 - 0.08) |
| Ribavirin | 40.52 | (49.96 - 32.7) | 1438.05 | (145378.48 - 14.22) | 442.08 | (494.52 - 395.21) | 35.15 | (49.55 - 24.93) |
| Aurintricarboxylic acid | 1.54 | (1.99 - 1.09) | 2.37 | (4.4 - 0.34) | 2.52 | (4.31 - 0.74) | 8.53 | (12.48 - 4.58) |
| Evans Blue | 1.89 | (3.93 - 0.14) | 2.4 | (4.5 - 0.3) | 4.88 | (8.92 - 0.84) | 4.76 | (5.77 - 3.76) |
| Patulin | 11.8 | (39.93 - 16.33) | 10.54 | (32.14 - 11.05) | 18.09 | (37.08 - 0.91) | not tested | |
| Suramin | 37.47 | (43.12 - 31.82) | 22.61 | (34.1 - 11.12) | 47.31 | 62.37 - 32.24) | 67.46 | (114.64 - 20.28) |

**Table S3. Individual $IC_{50}$ values for each hit evaluated in the four different inhibition assays against HMPV infection**. The geometric mean of each $IC_{50}$ value was calculated from at least three independent experiments performed in triplicate and are shown as the geometric mean with their 95% CI ( *n* =3).

**Figure S1**

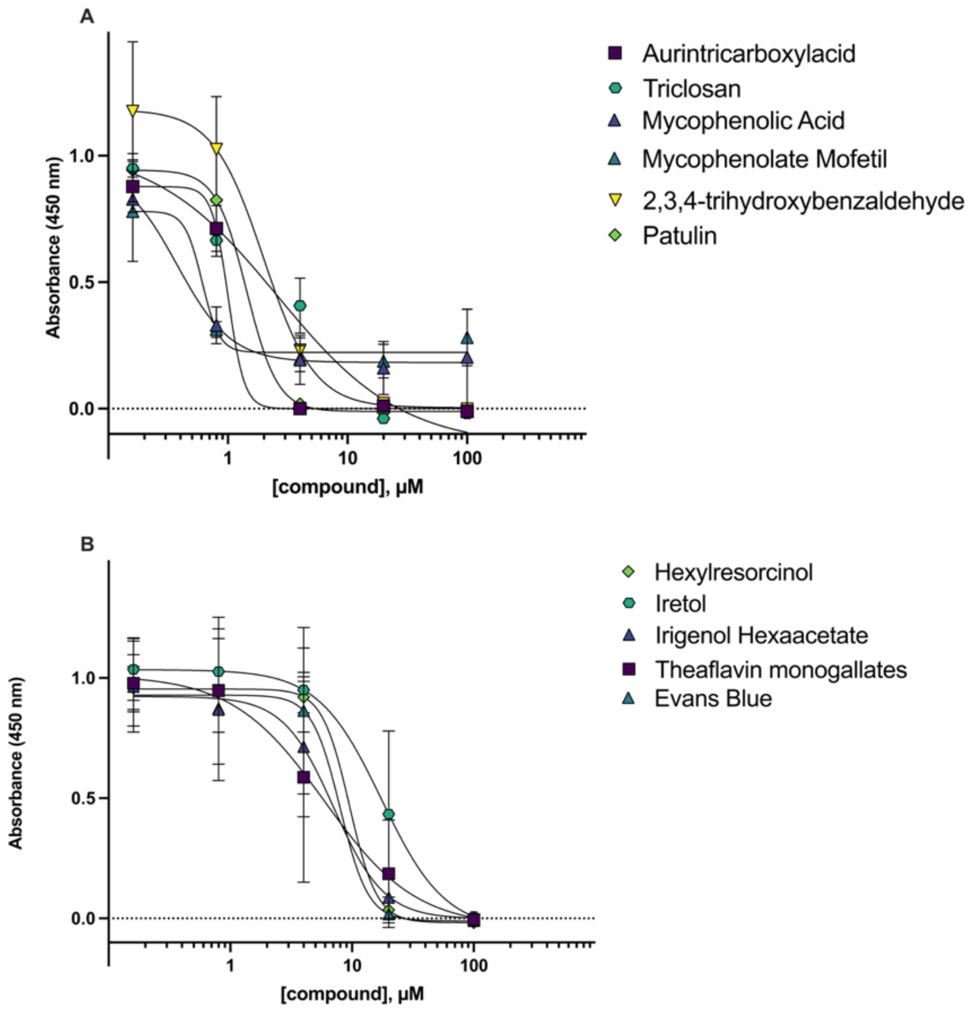

**Figure S1. Dose-dependent inhibition of identified drugs against HMPV-GFP infection of LLC-MK2 cells**. HMPV infectivity was measured by in situ ELISA (A) Drugs with $IC_{50}$ values below 10 µM. (B) Drugs with $IC_{50}$ values above 10 µM. Datapoints represent the mean value from two experimental repeats performed in technical duplicate, ± SD (*n* = 2).

**Figure S2**

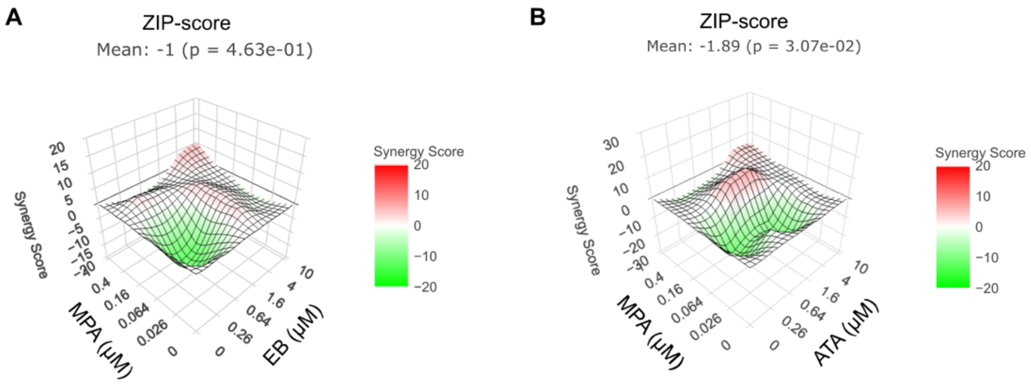

**Figure S2. Antiviral potency of drug combinations against HMPV *in vitro* infection using the ZIP method to determine synergy.** The calculations were performed using SyngerFinder (2) and the synergy score is visualised as the height of a 3D surface. (*n* = 1) (A) The ZIP-score and 3D surface of the synergy score of the combination of mycophenolic acid (MPA) with Evans Blue (EB). (B) The ZIP-score and 3D surface of the synergy score of the combination of MPA and aurintricarboxylic acid (ATA).
